# *Tritrichomonas muris* sensitizes the intestinal epithelium to doxorubicin-induced apoptosis

**DOI:** 10.1101/2024.08.08.607206

**Authors:** Nicolas V. Janto, Antoine R. Gleizes, Siyang Sun, Gurel Ari, Adam D. Gracz

**Affiliations:** Department of Medicine, Division of Digestive Diseases, Emory University; Graduate Program in Genetics and Molecular Biology, Emory University

**Keywords:** apoptosis, doxorubicin, intestinal epithelium, microbiome, Tritrichomonas muris

## Abstract

Doxorubicin (DXR) is a widely used chemotherapy drug that can induce severe intestinal mucositis. While the influence of gut bacteria on DXR-induced damage has been documented, the role of eukaryotic commensals remains unexplored. We discovered *Tritrichomonas muris* (*Tmu*) in one of our mouse colonies exhibiting abnormal tuft cell hyperplasia, prompting an investigation into its impact on DXR-induced intestinal injury. Mice from *Tmu*-colonized and *Tmu*-excluded facilities were injected with DXR, and tissue morphology and gene expression were evaluated at acute injury (6 h) and peak regeneration (120 h) phases. Contrary to previous reports, DXR did not significantly alter villus height, crypt depth, or crypt density in any mice. However, we did observe apoptosis, measured by cleaved caspase 3 (CC3) staining, in intestinal crypts at 6 h post-DXR that was significantly higher in mice colonized by *Tmu*. Interestingly, while DXR did not alter the expression of active and facultative intestinal stem cell (ISC) marker genes in control mice, it significantly reduced their expression in *Tmu*^+^ mice. *Tmu*, but not DXR, is also associated with increased inflammation and expression of the type 2 cytokines IL-5 and IL-13. However, pre-treatment of intestinal organoids with these cytokines is not sufficient to drive elevated DXR-induced apoptosis. These findings highlight the significant influence of commensal microbiota, particularly eukaryotic organisms like *Tmu*, on intestinal biology and response to chemotherapy, underscoring the complexity of gut microbiota interactions in drug-induced mucositis.

**NEW & NOTEWORTHY:** Our study found that the eukaryotic commensal *Tritrichomonas muris* (*Tmu*) significantly increases DXR-induced intestinal apoptosis in mice, despite no changes in tissue morphology. *Tmu* also reduces intestinal stem cell gene expression post-DXR injury, and elevates inflammation and type 2 cytokine expression in the absence of injury. *In vitro* organoid assays suggest that type 2 cytokines alone are insufficient to promote increased DXR-associated apoptosis. These findings emphasize the complex role of gut microbiota in drug-induced intestinal damage.

## INTRODUCTION

Doxorubicin (DXR) is a common chemotherapeutic with known gastrointestinal (GI) side effects, some of which can be severe enough to limit therapeutic application (1, 2). These side effects can be attributed to mucositis caused by intestinal epithelial cell (IEC) apoptosis, which has been well-documented in patients receiving chemotherapy and in rodent models treated with DXR (3–5). While the intestinal response to DXR has been well-characterized, evidence suggests that resident gut bacteria influence the type and extent of damage induced (6, 7). However, the gut microbiome also consists of viruses, fungi, and eukaryotes, which have not been considered for their potential impact on DXR-induced damage to the intestine.

Trichomonads are an order of flagellated eukaryotic protists generally considered to be commensal pathobionts. In recent years, several research groups have discovered trichomonads in the GI tracts of their mouse colonies consistent with *Tritrichomonas muris* (*Tmu*) (8–10). Since *Tmu* is considered to be non-pathogenic, few research animal facilities test for or intentionally exclude it (11). Despite its non-pathogenic classification, *Tmu* and related protozoa have been shown to significantly alter intestinal biology and phenotypes, primarily through their influence on the mucosal immune landscape (8–10, 12–14).

Several research groups incidentally discovered *Tmu* in their mice after noticing abnormally elevated tuft cell frequencies in the intestinal epithelium (9, 10). *Tmu* and related protozoa have been shown to drive tuft cell hyperplasia in the small intestine through a type 2 innate immune response. Chemosensory receptors on tuft cells detect succinate produced by the trichomonad and activate a tuft cell-ILC2 circuit that promotes further tuft cell differentiation (9, 13, 15). In addition to activation of type 2 immunity, Th1 and Th17 T cell proliferation as well as inflammasome activation have been observed in the GI tracts of mice colonized with trichomonads (8, 12, 14). With evidence of *Tmu* colonization in humans, it is possible that *Tmu* and similar eukaryotes impact a wide range of pathophysiology in the human intestine, including inflammatory responses to mucosal injury (14).

Th1, Th2, and Th17 cytokines have been shown to regulate intestinal stem cell (ISC) renewal, differentiation, proliferation, and gene expression (16–18). Additionally, tuft cells are implicated in epithelial repair and regeneration following injury of the small intestine (19, 20). Following the incidental discovery of *Tmu* in one of our mouse colonies, we decided to study its impact on intestinal epithelial injury. Since both the tuft cell and mucosal T cell populations are altered in the presence of *Tmu*, we hypothesized that *Tmu* colonization would influence the response to DXR-induced injury. Here, we find that *Tmu*-colonized mice exhibit higher rates of IEC apoptosis, reduced proliferation, and greater loss of ISCs in response to DXR.

## MATERIALS AND METHODS

### Mice

All experiments were carried out on wild type C57Bl/6 adult males 10-20 weeks of age, except for experiments collecting feces for Lipocalin-2 quantification which were carried out on wild type C57Bl/6 males and females 10-25 weeks of age. All mice received PicoLab Rodent Diet 20 (LabDiet, 5053) and water *ad libitum.* The Institutional Animal Care and Use Committee (IACUC) of Emory University reviewed and approved all animal protocols.

### *Tmu* detection by bright field microscopy

*Tmu* colonization status of all mice was confirmed as previously described (21). Briefly, the cecum was dissected, cut open longitudinally, and placed in a 50 mL conical containing 10 mL Dulbecco’s Phosphate-Buffered Saline (DPBS) (Gibco, 14190250). The conical was shaken vigorously to release all cecal contents. The contents were then diluted by bringing the volume up to 50 mL with DPBS. 10 μL aliquots were examined by bright field microscopy. *Tmu*-colonized samples exhibited abundant highly motile teardrop shaped flagellated protists, which were entirely absent from uninfected samples.

### Stool DNA extraction and *Tmu* validation by PCR

DNA was extracted from fecal samples per the manufacturer’s protocol for the Quick-DNA Fecal/Soil Microbe MiniPrep Kit (Zymo, D6010). *Tritrichomonas muris* was detected by PCR with previously published primers recognizing *Tmu* 28S rRNA gene: 5′-GCTTTTGCAAGCTAGGTCCC-3′ and 5′-TTTCTGATGGGGCGTACCAC-3′ (9).

### Doxorubicin injection

Mice received one intraperitoneal injection of 20 mg/kg doxorubicin HCl (Adryamicin, Selleck Chemicals, S1208). Mice were monitored for weight loss daily. Any mouse exhibiting 20% or greater weight loss following doxorubicin administration was humanely euthanized, per IACUC guidelines. Untreated control mice did not receive a vehicle injection except for experiments in which fecal samples were collected daily for Lipocalin detection, which were injected with an equal volume of Dulbecco’s Phosphate-Buffered Saline (DPBS) (Gibco, 14190250) to control for any inflammatory effects of the injection itself.

### Hematoxylin and eosin (H&E) staining

Duodenum was dissected, flushed with 1X PBS, fixed overnight in 4% PFA at 4°C, then transferred to 100% EtOH. H&E staining was performed by the Cancer Tissue and Pathology Shared Resource of Winship Cancer Institute of Emory University on a Sakura Tissue-Tek Prisma Automated Slide Stainer and Coverslipper (Sakura Finetek) and the Tissue-Tek Prisma H&E Stain Kit #1 (Sakura Finetek, 6190).

### Immunofluorescent staining

Jejunum was dissected, flushed with 1X PBS, fixed overnight in 4% PFA at 4°C, then transferred to 30% sucrose for an additional 16-18 h prior to embedding in Tissue-Plus OCT (Fisher Scientific; 23-730-571) and freezing at -70°C. Immunofluorescence was carried out on 10 μm sections, cut on a cryostat (Leica; CM1860). Tissue was rehydrated with PBS and then permeabilized with 0.3% Triton X-100 (Sigma; T8787-250ML) in PBS for 20 min at room temperature, followed by blocking in 5% normal donkey serum (NDS; Jackson Immuno; 017-000-121) for 45 min at room temperature. Primary antibodies were diluted in 1X PBS and incubated at 4°C overnight. Secondary antibodies were diluted in 1X PBS and incubated for 45 min at room temperature. Nuclear counterstain was carried out using DAPI (Millipore Sigma; D9542) diluted 1:2000 in PBS. Primary antibodies were used at the following concentrations for immunostaining: 1:200 anti-CD326 (EPCAM; BioLegend; 118201), 1:100 anti-DCAMK1 (DCLK1; Abcam; ab192980), 1:400 anti-Cleaved caspase 3 (CC3; Cell Signaling; 9661S), 1:200 anti-OLFM4 (Cell Signaling; 39141S). Secondary antibodies used for immunostaining: 1:250 donkey anti-rabbit Alexa Fluor 555 (ThermoFisher; A31572), 1:250 donkey anti-rat Alexa Fluor 488 (ThermoFisher; A21208).

### EdU staining

Mice received an intraperitoneal injection of 500 μg of EdU (Click Chemistry Tools; 1149-500) two hours prior to sacrifice. Staining for EdU was carried out on 10 μm section with the Click-&-Go Plus 555 Imaging Kit (Vector Laboratories, Inc.; CCT-1317) per the manufacturer’s protocol.

### Image acquisition and quantification

Representative images were acquired on a laser scanning Nikon A1R HD25 Confocal Microscope in the Emory Integrated Cellular Imaging Core. Images for quantification were taken using an Olympus IX-83 inverted epifluorescent microscope and cellSens Imaging software. For tissue sections, 30 crypts and/or villi were quantified per biological replicate. For organoids, 90 organoids were quantified per experimental group.

### RNA isolation and RT-qPCR

A portion of jejunum < 5 mm in length was placed in 200 µL of RNAlater stabilization solution (Thermo Fisher, AM7020) and stored at room temperature for up to a week. 5-8 mg of tissue were then transferred to 500 µL of RNA Lysis Buffer from the Ambion RNAqueous Micro kit (Thermo Fisher, AM1931), in which it was homogenized using a Bio-Gen PRO200 Homogenizer (Pro Scientific, 01-01200). Total RNA was purified using the Ambion RNAqueous Micro kit according to manufacturer instructions. All samples were treated with DNase (Ambion) for 30 min at 37°C and DNase inactivated following manufacturer protocol. Total RNA was quantified using a Qubit v3 Fluorometer (Thermo Fisher; Q33216) and the Qubit High Sensitivity RNA Quantification Assay (Thermo Fisher; Q32855). 100 ng of RNA was subjected to reverse transcription using the iScript cDNA Synthesis Kit (Bio-Rad; 1708891) and diluted 1:5 in molecular grade water (Corning; 46-000-CI). RT-qPCR was carried out in technical triplicate using Taqman assays and SsoAdvanced Universal Probes Supermix (Bio-Rad; 1725284), following manufacturer protocols and using 1 uL diluted cDNA input per 10 uL reaction. Reactions were run on a QuantStudio 3 Real Time PCR instrument (Thermo Fisher) and analyzed using the ΔΔC_T_ method (22). 18S was selected as the internal housekeeping gene. Taqman assay IDs used in this manuscript are: *18S* (Hs99999901_s1), *Ascl2* (Mm01268891_g1), *Ccl11* (Mm00441238_m1), *Clu* (Mm01197001_m1), *Il13* (Mm00434204_m1), *Il25* (Mm00499822_m1), *Il33* (Mm00505403), *Il4* (Mm00445259_m1), *Il5* (Mm00439646_m1), *Lgr5* (Mm00438890_m1), *Ly6a* (Mm00726565_s1), *Pou2f3* (Mm00478293_m1), *Sox4* (Mm00486320_s1).

### Lipocalin-2 quantification

Mice were injected with DXR or DPBS as described above (*see **Doxorubicin injection***). Mouse feces were harvested in a 1.5 mL conical tube the day of injection and every day for 5 days after. Samples were either frozen at -20°C or processed fresh. Feces were weighed on a microscale and resuspended in 1x PBS 0.1% Tween-20 at 100 mg/mL. Samples were then vortexed for 20 min at room temperature and centrifuged for 10 min at 12,000 g at 4°C. Supernatant was harvested and total protein concentration was quantified using Pierce^TM^ BCA Protein Assay Kit (Thermo Scientific, 23225) according to manufacturer’s instructions. Each sample was diluted down to 500 μg/mL of total protein in 1x PBS before performing Lipocalin-2 titration to maintain a range appropriate to the quantification assay. Lipocalin-2 titration was performed using the DuoSet^©^ ELISA Mouse Lipocalin-2/NGAL kit (R&D Systems, DY1857-05) according to manufacturer’s instructions.

### Crypt isolation

Crypts were isolated for organoid culture as previously described(23). Briefly, intestines were dissected out, the first 6cm discarded, and the remaining proximal half designated as jejunum and taken for organoid isolation. Jejunal segments were opened longitudinally and rinsed briefly in a 50 mL conical containing 10mL sterile Dulbecco’s Phosphate-Buffered Saline (DPBS) (Gibco; 14190250). Tissue was transferred to a new 50mL conical containing 5 mM EDTA (Corning; 46-034-CI) in 10 mL sterile DPBS and incubated at 4°C on a rocking platform set to 30 RPM for 12 min. Tissue was retrieved and villi removed by gentle “brushing” with a P200 pipette tip on a glass plate. Intestinal tissue was then rinsed briefly in a petri dish containing sterile DPBS and cut into pieces 2-3 cm in length before being transferred to a new 50 mL conical containing 5 mM EDTA in 10 mL sterile DPBS and incubated at 4°C on a rocking platform for 35min. Jejunal pieces were transferred to a 50 mL conical containing 10 mL sterile DPBS and shaken for 3-4 min to release crypts, confirming expected crypt density and morphology by light microscopy at 1min intervals to determine when to end the dissociation protocol. 10 mL sterile DPBS was added to isolated crypts, which were then filtered through a 70 μm cell strainer, pelleted at 600 g for 5 min at RT, and resuspended in 200-500 uL Advanced DMEM/F12 (Gibco; 12634010).

### Organoid culture

Crypt density per 10uL media was examined qualitatively by light microscopy and crypt density estimated, with the goal of determining volume required to achieve ∼300 crypts per 30 μL Matrigel in culture. Crypts were resuspended in 80% phenol red-free, growth factor-reduced Matrigel (Corning; 356231) and plated as 30 μL droplets in 48-well plates. Matrigel was allowed to polymerize at RT for 5 min, then at 37°C for 15 min. ENR media was overlaid at 200 μL per well: Advanced DMEM/F12, 1X N2 (Thermo Fisher; 17502048), 1X B27 w/o vitamin A (Thermo Fisher; 12587010), 1X HEPES (Gibco; 15630080), 1X Penicillin/Streptomycin (Sigma-Aldrich; P4333-100ML), 1X Glutamax (Gibco; 35050061), 10% RSPO1-CM (made using RSPO1 transfected HEK293T cells following manufacturer protocol; Sigma Aldrich; SCC111), 50ng/mLrecombinant murine EGF (Gibco; PMG8041), and 100ng/mL recombinant human NOGGIN (PeproTech; 120-10C-20UG). 500μg/mL Primocin (Invitrogen; ant-pm-1) and 10mM Y27632 (Selleck Chemicals; S1049) were added to overlay media for the first 48hr after plating crypts and then excluded from all other media changes. ENR media was replaced every 48 hours.

### Organoid treatment

After allowing organoids to establish for 48 hours, either 100 ng/mL recombinant mouse IL-5 (Biolegend, 581502) or 100 ng/mL recombinant mouse IL-13 (Biolegend, 575902) was added to each organoid media change for the remainder of the experiment, for a total of 4 days of cytokine treatment. Six days after the organoids were plated, 0.5 μg/mL doxorubicin (Selleck Chemicals, S1208) was added to the media for 4 hours, after which time organoids were processed immediately for whole-mount staining (*see **Organoid recovery and whole-mount staining***).

### Organoid recovery and whole-mount staining

Organoids were recovered and stained as previously described (24). Briefly, organoids were rinsed with 1 mL PBS and then recovered from the Matrigel matrix by replacing the culture media with 1 mL cold Cell Recovery Solution (Corning, 354253) and incubating at 4°C for 45 minutes. The organoids were then transferred to 15 mL conical tubes pre-coated with 1% BSA (Sigma-Aldrich, A9647) in PBS. The tubes were filled to 10 mL with PBS and spun down at 70 g for 3 min at 4°C. The organoids were resuspended in 1 mL 4% PFA and incubated at 4°C for 45 min. The tube was then filled to 10 mL with cold 0.1% (vol/vol) PBS-Tween (PBT) and incubated for 10 min at 4°C before spinning down at 70g for 5 min at 4°C. The organoids were resuspended in 1 mL cold organoid washing buffer (OWB; 1 mL of Triton X-100 and 2 g of BSA in 1 liter of PBS), transferred to a low-adherence 96-well plate, and incubated at 4°C for 15 min. Then organoids were incubated overnight at 4°C in 1:400 anti-Cleaved caspase 3 (CC3; Cell Signaling; 9661S). Organoids were rinsed 3 x 2 hours in OWB on a rocking platform before incubating overnight at 4°C in 1:250 donkey anti-rabbit Alexa Fluor 555 (ThermoFisher; A31572). Organoids were rinsed for 1 hour in OWB before incubating at room temperature for 20 minutes in 1:2000 DAPI (Millipore Sigma; D9542). Organoids were again rinsed 3 x 2 hours in OWB, then transferred to 1.5 mL tubes and pelleted at 70 g for 2 min at 4°C. Organoids were resuspended in 60% vol/vol glycerol and 2.5 M fructose and incubated at room temperature for 20 min. To facilitate imaging, organoids were mounted one a glass slide with a cover slip sitting atop two layers of double-sided tape, as previously described (24). All wash steps and antibody incubations were carried out on a rocking platform.

### Statistical analysis

Statistical analyses were carried out in Prism 10.2.2 (GraphPad Software). For statistical comparison, unpaired t test was used for datasets of two groups, ordinary one-way ANOVA was used for datasets of four groups, and two-way ANOVA was used for datasets of six groups. Pairwise comparisons were carried out using a Šídák’s multiple comparisons test. A value of p < 0.05 was considered significant. All values are depicted as mean ± SEM. For ease of visual presentation, the results of full pairwise comparisons are shown in Supplemental Table S1.

## RESULTS

### Facility-dependent tuft cell hyperplasia leads to *Tmu* detection

While conducting routine histology experiments, we noticed a difference in tuft cell frequency in mice from colonies housed in two different facilities at Emory University, HSRB and WBRB (Fig. 1A and B). Since HSRB is a barrier facility that specifically excludes protists, we hypothesized that *Tmu* colonization might be responsible for WBRB-associated tuft cell hyperplasia, as previously reported (9). Upon examination of cecal contents, mice housed in WBRB showed evidence of unicellular flagellates whose size and morphology are consistent with *Tmu,* while mice house in HSRB did not (Fig. 1D). PCR amplification of purified genomic DNA from the fecal contents of our mice using primers designed to specifically detect the 28S rRNA gene of *Tmu* validated their identity and facility-specific colonization (Fig. 1C) (9). Together these data demonstrate that *Tmu* is present in the WBRB colony displaying tuft cell hyperplasia, but not in the HSRB barrier facility mouse colony.

**Figure 1:**
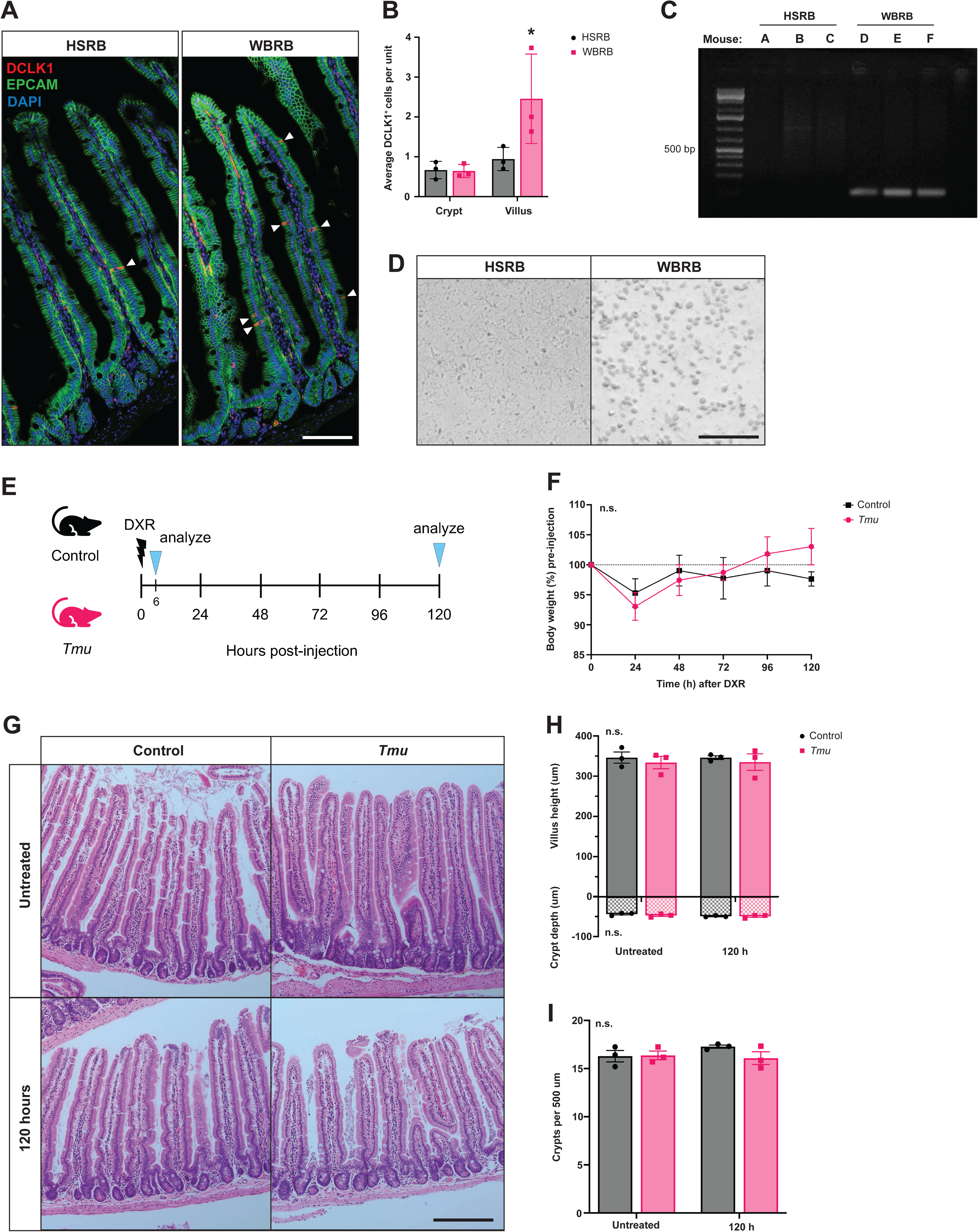
No change in intestinal morphology in response to DXR regardless of *Tmu*. (A) Immunofluorescent staining for DCLK1 reveals facility-dependent tuft cell hyperplasia (scale bar represents 100 µm). (B) DCLK1^+^ tuft cell frequency per villus is significantly greater in mice housed in WBRB, while no difference in tuft cell frequency is seen in the crypts (n = 3 replicates per facility, p < .001). (C) Cecal contents from mice housed in each facility reveal that a unicellular flagellate is abundantly present in mice housed in WBRB and absent from mice housed in HSRB (scale bar represents 100 µm). (D) PCR amplification of the *Tmu* 28S rRNA gene from purified fecal DNA collected from mice residing in both facilities indicates that *Tmu* is present in WBRB mice, and absent in HSRB mice. (E) Schematic of experimental design, where mice either uninfected or colonized with *Tmu* are injected with DXR and tissue is collected at 6 hours and 120 hours post-injection. (F) H&E staining of duodenum reveals no readily observable difference in morphology between DXR-treated and untreated controls or between *Tmu^+^* and *Tmu^-^* samples (scale bar represents 200µm). (G) No changes in crypt height or crypt depth in response to DXR or *Tmu* (n = 3 replicates per group, n.s. indicates not significant). (H) Crypt density measured over 500 um intervals does not change in response to DXR or *Tmu* (n = 3 replicates per group, n.s. indicates not significant). (I) Body weight as percent of starting body weight recorded daily for 5 days following DXR injection reveals no significant change in body weight over the course of the experiment or between *Tmu* mice and controls (n = 3 replicates per group).

### *Tmu* does not impact intestinal morphological response to DXR

Because we were carrying out unrelated studies using DXR as a damage model in both HSRB and WBRB, we next sought to determine if the presence of *Tmu* might be driving phenotypic differences in damage response between the two facilities. To investigate the impact of *Tmu* on the intestinal response to DXR, we gave a single IP injection of 20 mg/kg to mice from both facilities (Fig. 1E). Mice were weighed daily, revealing no significant difference in body weight response to DXR between *Tmu*^+^ mice and controls (Fig. 1F). The presence or absence of *Tmu* was validated by examination of cecal contents post-sacrifice, which confirmed facility-specific colonization. Previous studies have reported extensive morphological changes in the intestinal crypt-villus axis following acute DXR injury and dependent on the presence of the microbiome (4, 7). To examine the potential impact of *Tmu* on these changes, we collected tissue at 120 h post-DXR, which has been reported to be a “peak” regenerative timepoint with distinct morphological phenotypes including villus blunting, crypt hyperplasia, and crypt loss (4, 7). To evaluate crypt and villus morphology we performed H&E staining. However, we saw no appreciable differences in villus height or crypt density at 120 h post-DXR compared to untreated controls (Fig. 1G-I). It also appears that *Tmu* does not impact these morphological metrics in the absence of cytotoxic injury. These data suggest that changes in body weight and gross intestinal morphology are not reliable indicators of DXR-induced injury or *Tmu* colonization.

### *Tmu*-colonized mice exhibit elevated rates of DXR-induced apoptosis in vivo

We next looked to apoptosis as a measure of DXR-induced damage and collected tissue at 6 h post-DXR, which was previously reported to represent peak cell death (4). We stained for cleaved caspase 3 (CC3) and found that *Tmu* alone does not influence apoptosis, but that *Tmu*-colonized mice exhibited significantly more apoptosis at 6 h post-DXR than those lacking *Tmu* (Fig. 2A and B). Since DXR preferentially induces apoptosis in ISCs and early progenitors, we wondered if the increased apoptosis observed in *Tmu*^+^ mice was being driven by increased sensitivity of a specific ISC/progenitor compartment (4). For each crypt, we demarcated a boundary separating the ISC compartment (below) from the transit amplifying (TA) progenitors (above), based on the location of Paneth cells (Fig. 2C). We then calculated the percentage of CC3^+^ cells above and below that boundary, revealing no change in bias of DXR’s target in the presence of *Tmu*. Taken together, these results reveal an increase in sensitivity to DXR-induced apoptosis in mice colonized with *Tmu* that is not biased toward the ISC or TA compartment.

**Figure 2:**
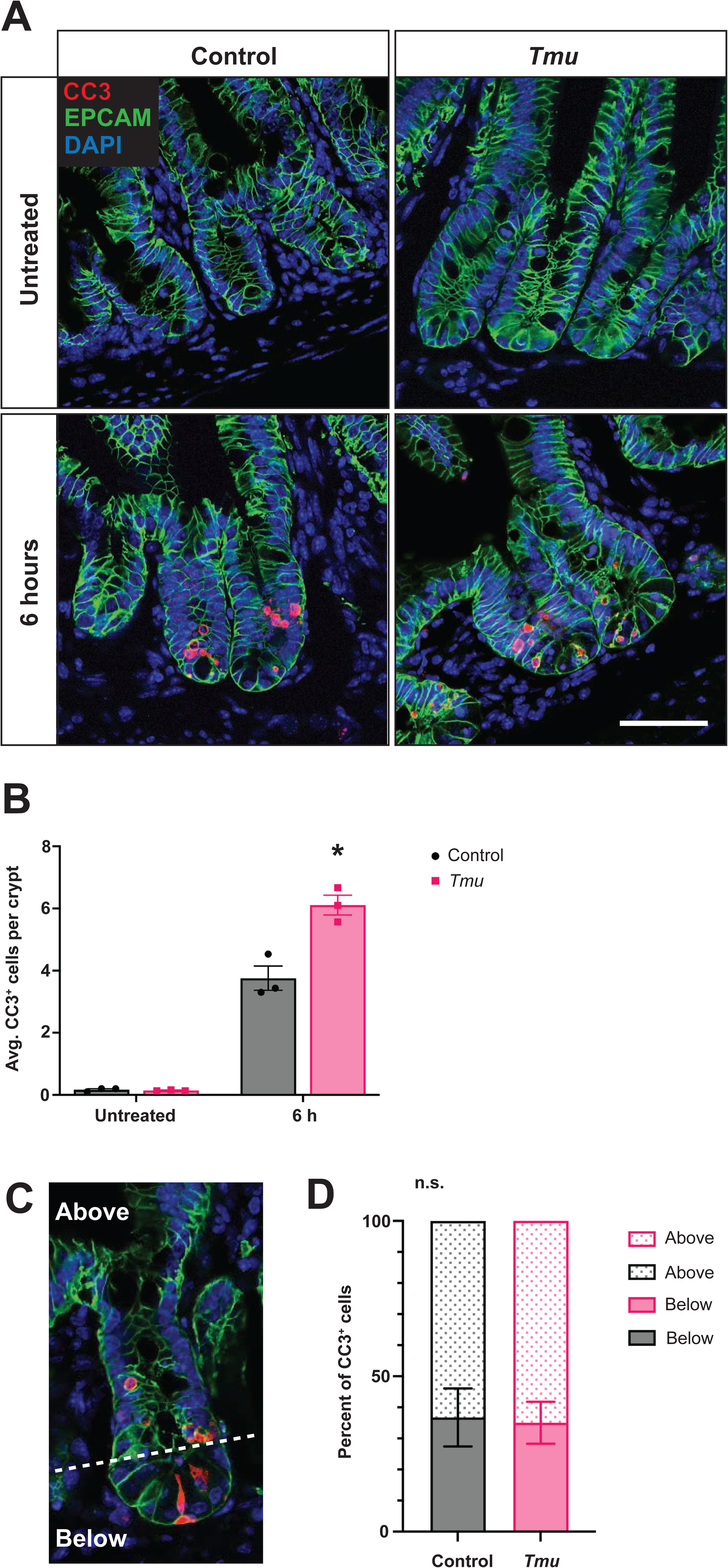
Acute apoptosis in response to DXR is increased in *Tmu*^+^ mice. (A) Immunofluorescent staining for CC3 in jejunum of DXR-treated and untreated mice both with and without *Tmu* (scale bar represents 50 µm). (B) Frequency of CC3^+^ cells per crypt increases in response to DXR in both sets of mice, but is significantly higher in mice colonized with *Tmu* (n = 3 replicates per group, p < 0.01). (C) Representative boundary separating ISC and TA compartments. (D) Percent of CC3^+^ cells in the crypt at 6 h post-DXR that fall either above or below the boundary shown in (C) (n = 3 replicates per group, n.s. indicates not significant).

### Presence of *Tmu* alters ISC response to DXR

To assess the ISC response following DXR treatment, we looked at crypt cell proliferation by EdU as well as ISC gene expression. Both *Tmu*^+^ mice and *Tmu*^-^ control crypts demonstrate a significant reduction in proliferation at 6 h post-DXR compared to their untreated controls (Table S1). However, *Tmu*^+^ mice exhibited significantly less proliferation than *Tmu*^-^ controls at 6 h post-DXR, with no significant differences between groups at 120 h or in the absence of DXR (Fig. 3A-B). We also stained for OLFM4, an ISC marker which has been shown to drastically diminish in expression following DXR treatment (6, 25). While we did observe an initial downward trend in OLFM4 staining at 6 h post-DXR that was significant in *Tmu*^+^ mice, it was followed by a significant increase in OLFM4 from 6 h to 120 h (Supplemental Table S1). OLFM4 staining recovered to a significantly greater extent in *Tmu*^-^ controls at 120 h, resulting in significantly more OLFM4^+^ cells per crypt than there were in an uninjured state (Fig. 3C-D, Supplemental Table S1).

**Figure 3:**
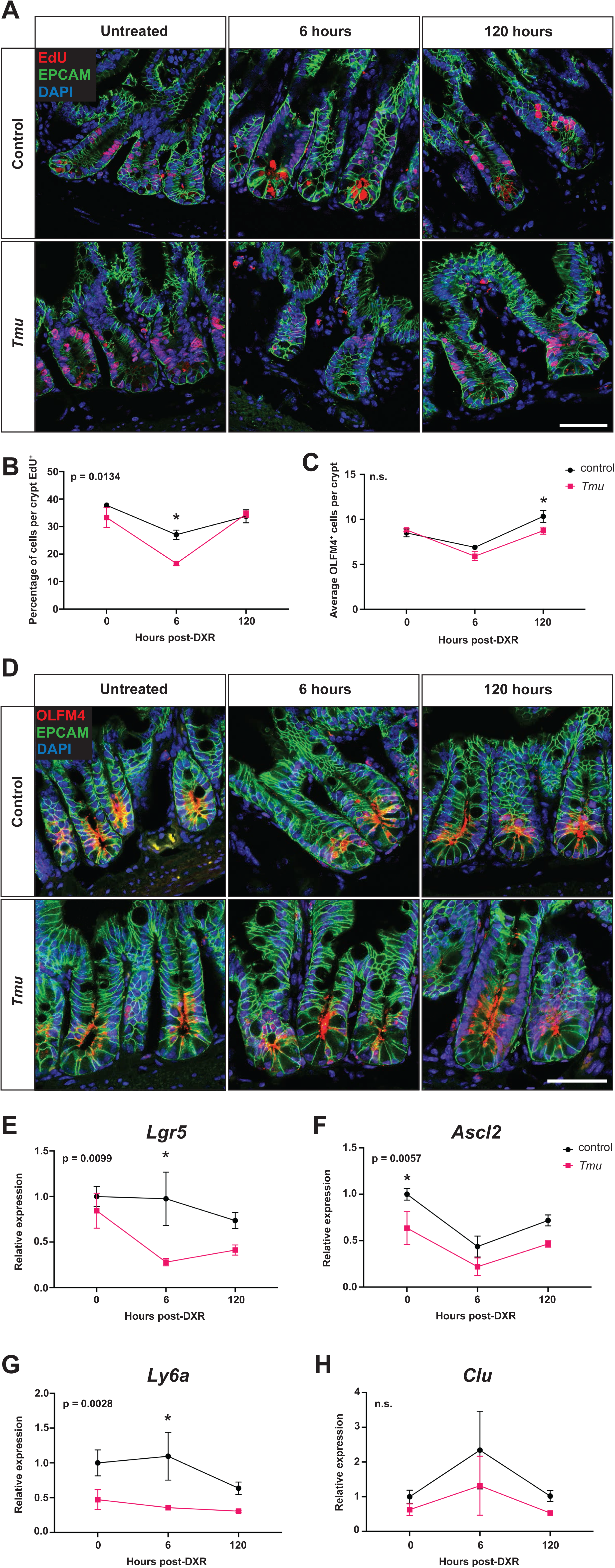
Altered ISC response to DXR in the presence of *Tmu*. (A) Staining for EdU following a 2 h EdU pulse shows a reduction in proliferation at 6 h post-DXR (scale bar represents 50 µm). (B) *Tmu^+^* mice exhibit a significant reduction in the proportion of EdU^+^ cells per crypt (n = 3 replicates per group, listed p-value indicates significant difference between lines, * indicates significant difference at a particular timepoint where p < 0.01). (C) No significant difference in frequency of OLFM4^+^ ISCs per crypt is seen in response to DXR or *Tmu* (n = 3 replicates per group, n.s. indicates not significant). (D) Representative images of immunofluorescent staining for OLFM4 (scale bar represents 50 µm). RT-qPCR of aISC genes (E) *Lgr5* and (F) *Ascl2* as well as fISC genes (G) *Ly6a* and (H) *Clu* reveal a more pronounced loss of *Lgr5* and *Ly6a* expression at 6 h post-DXR and lower baseline expression of *Ascl2* and *Ly6a* in *Tmu*^+^ samples (n = 3 replicates per group, listed p-value refers to the significance of *Tmu*’s contribution to variation in the data, * indicates significant difference between *Tmu* status groups at the same timepoint, p < 0.05).

In addition to loss of OLFM4, DXR is associated with loss of other markers of active ISCs (aISCs), including *Lgr5*, while *Ly6a* and *Clu* are both known to be upregulated during repair in facultative ISCs (fISCs) (6, 25–28). We assessed aISC (Fig. 3E-F) and fISC (Fig. 3G-H) gene expression by RT-qPCR on RNA purified from whole intestinal tissue. Although *Lgr5* has been shown to progressively decrease over the first 120 h following DXR treatment, we only saw a downward trend in *Lgr5* expression following DXR in *Tmu*^+^ samples, where *Lgr5* expression was significantly lower in *Tmu*^+^ samples at 6 h post-DXR compared to *Tmu*^-^ controls (6) (Fig. 3E). We did see a significant reduction in expression of the aISC-promoting transcription factor *Ascl2* in all samples at 6 h post-DXR, however *Tmu*^+^ samples displayed significantly lower expression prior to injury and trended lower at both post-injury timepoints (Fig. 3F, Supplemental Table S1). Curiously, neither *Ly6a* nor *Clu* significantly increased in expression in response to DXR in either the *Tmu*^+^ mice or the controls, though *Clu* exhibited an upward trend at 6 h post-injection (Fig. 3G-H, Supplemental Table S1). We do, however, see that *Ly6a* expression is significantly lower in *Tmu*^+^ samples at 6 h post-DXR (Fig. 3G). Overall, these results suggest that some aISC and fISC gene expression is lower in the presence of *Tmu*, in the absence of and following injury, and that *Tmu* colonization is associated with a greater reduction in proliferation following DXR injury.

### Tuft cell response to DXR-induced injury is modulated by *Tmu*

Given the influence of *Tmu* colonization on tuft cell numbers as well as the involvement of tuft cells in intestinal regeneration, we decided to look at DCLK1 expression in the intestinal epithelium at both post-DXR timepoints to see how tuft cells respond to DXR in the of *Tmu* (19, 20). Previous studies report a loss of tuft cells in response to DXR, which is consistent with trends observed in our *Tmu*^-^ tissue (Fig 4A-C) (4). However, we see the opposite initial effect in our *Tmu*^+^ samples, where DCLK1 expression increases at 6 h post-injection before dropping below baseline levels by 120 h (Fig. 4A-C). While a change in cell populations due to differentiation would be unlikely within 6 h of DXR treatment, we hypothesized that the changes in DCLK1 protein expression may be driven by an increase in expression of pro-tuft cell transcription factors. To this end, we measured expression of *Pou2f3* and *Sox4* by RT-qPCR, revealing that neither transcription factor increases in expression in response to DXR (Fig. 4D-E). These data suggest that the significant DXR-induced changes in DCLK1 expression in *Tmu*^+^ tissue, are not driven by gene expression changes of canonical tuft cell transcription factors.

**Figure 4:**
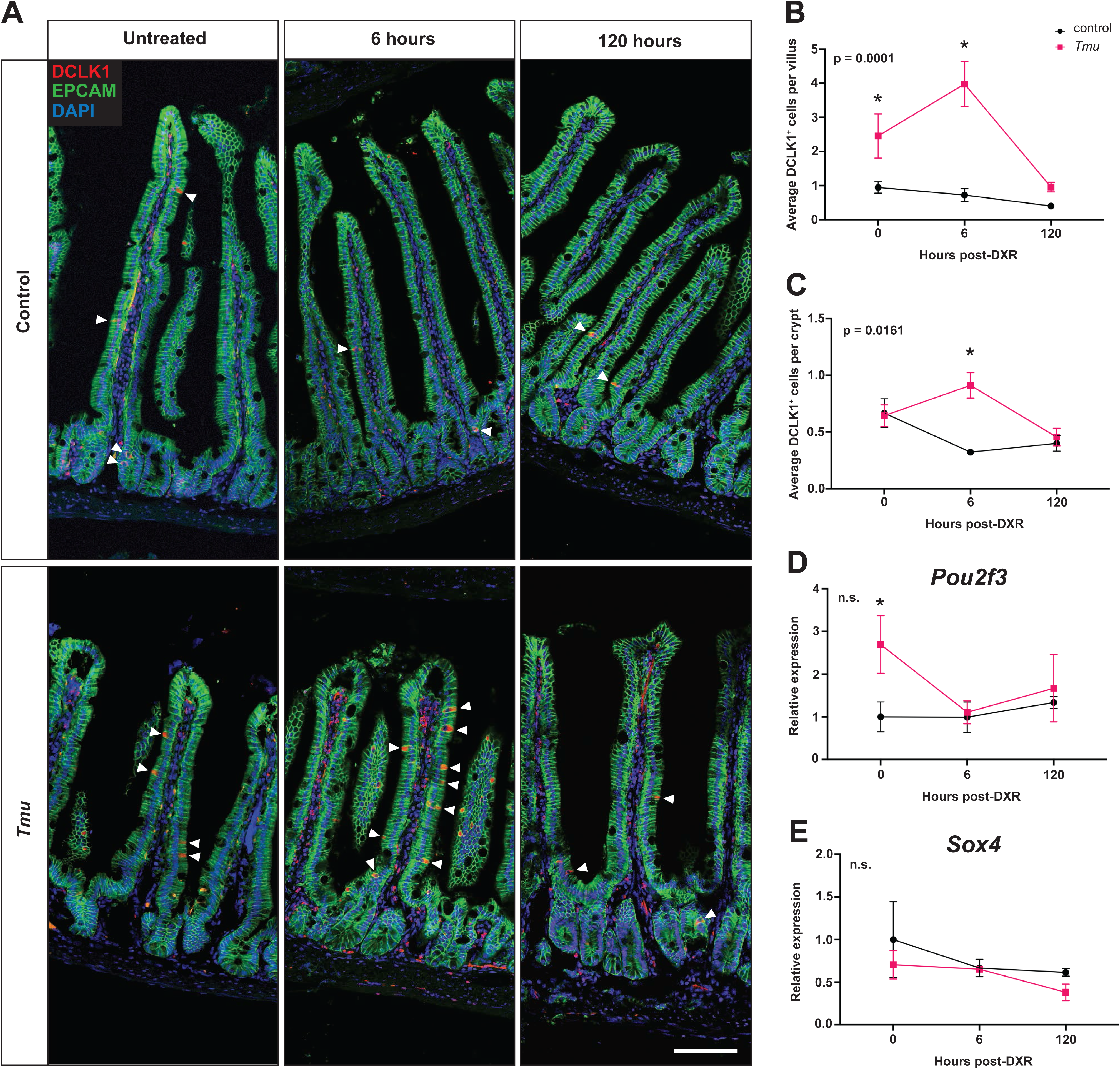
*Tmu* changes tuft cell response to DXR independent of transcription factor gene expression. (A) Immunofluorescent staining of DCLK1 demonstrates altered tuft cell response to DXR in the presence of *Tmu* (scale bar represents 100 µm). DCLK1^+^ cell frequencies trend downward in response to DXR in *Tmu^-^* controls, but increase before dropping below baseline in *Tmu*^+^ mice in both (B) villi and (C) crypts (n = 3 replicates per group, listed p-value refers to the significance of *Tmu*’s contribution to variation in the data, * indicates significant difference between *Tmu* status groups at the same timepoint, p < 0.01). RT-qPCR of pro-tuft cell transcription factors, (D) *Pou2f3* and (E) *Sox4*, reveals no significant change in expression in response to DXR (n = 3 replicates per group, n.s. indicates not significant).

### *Tmu*, not DXR, is associated with inflammation and type 2 cytokine expression

Since *Tmu* is known to drive tuft cell hyperplasia through a type 2 innate immune response, we sought to characterize the impact of DXR on the immune and inflammatory landscape in the small intestine of mice colonized with *Tmu*. We originally began collecting fecal samples from mice to measure the amount of Lipocalin-2 (Lcn-2), a biomarker of intestinal inflammation produced by neutrophils and IECs that’s upregulated in response to damage, as a means of monitoring inflammation over the course of DXR treatment (29). Surprisingly, DXR had no significant effect on Lcn-2 production in mice housed in either facility, and significant differences in Lcn-2 production could only be explained by the presence of *Tmu*, as determined by a three-way ANOVA (p < 0.0001) (Fig. 5A).

**Figure 5:**
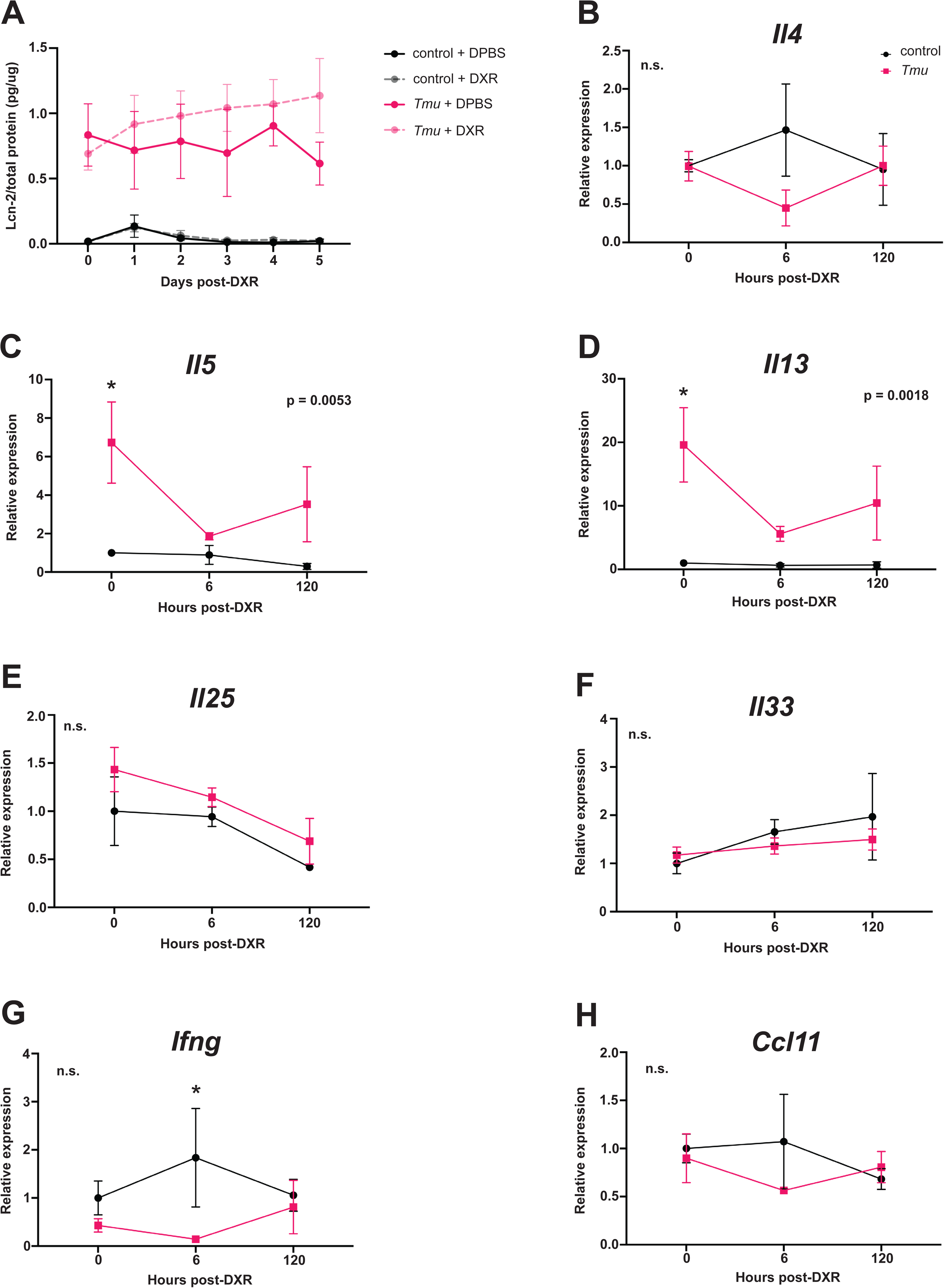
Elevated inflammation and type 2 cytokines in *Tmu*-colonized mice. (A) Amount of Lipocalin-2 (Lcn-2) per μg total protein as detected by ELISA reveal that Lcn-2 expression is driven more by *Tmu* than by DXR (n = 4 for control + DPBS, n = 7 for control + DXR, n = 3 for *Tmu* + DPBS, and n = 5 for *Tmu* + DXR). Expression of type 2 cytokines (B) *Il4*, (C) *Il5*, and (D) *Il13* as determined by RT-qPCR support that an active type 2 immune response is present in *Tmu*^+^ mice, characterized by significantly elevated *Il5* and *Il13* expression that drops after DXR treatment (n = 3 replicates per group, listed p-value refers to the significance of *Tmu*’s contribution to variation in the data, * indicates significant difference between *Tmu* status groups at the same timepoint, p < 0.01, n.s. indicates not significant). Expression of IEC-produced cytokines (E) *Il25* and (F) *Il33* is not significantly different in *Tmu*^+^ mice compared to uninfected controls (n = 3 replicates per group, n.s. indicates not significant). *Tmu*^+^ mice also do not exhibit significantly different expression of (G) *Ifng* or (H) *Ccl11* compared to uninfected controls (n = 3 replicates per group, n.s. indicates not significant).

To characterize the immune landscape, we performed RT-qPCR on RNA purified from whole jejunum, starting with known ILC2-secreted cytokines IL-4, IL-5, and IL-13 (30). *Il4* was not elevated in *Tmu*-colonized mice, but both *Il5* and *Il13* were elevated in the absence of DXR injury, indicating an active type 2 innate immune response (Fig 5B-D). Curiously, both *Il5* and *Il13* expression significantly declined at 6 h post-DXR (Fig. 5C-D, Supplemental Table S1). Also surprising was that neither *Il25* nor *Il33* expression were elevated in *Tmu*^+^ samples, despite being IEC-secreted cytokines that activate ILC2s (Fig. 5E-F). We also looked at expression of *Ifng*, associated with Th1 immune responses, and *Ccl11*, an eosinophilic attractant upregulated during parasitic infection and a regulator of intestinal inflammation (31, 32). Neither gene was significantly elevated in *Tmu*^+^ samples, confirming that the type 2 immune response is the dominant immune response to *Tmu* colonization (Fig. 5G-H). *Ifng* expression was significantly greater in *Tmu*^-^ controls compared to *Tmu*^+^ samples, however the transcript’s expression wasn’t significantly impacted over the course of DXR treatment in either group (Fig. 5G, Supplemental Table S1). Together, these results suggest that *Tmu* is strongly associated with an active type 2 immune response at homeostasis as well as inflammation, and that DXR does not significantly upregulate inflammatory markers or immune cytokine expression.

### Type 2 cytokines do not exacerbate DXR-induced apoptosis *in vitro*

Seeing that both *Il5* and *Il13* were significantly upregulated in our *Tmu*^+^ mice, we wanted to investigate whether these cytokines were driving the excess apoptosis observed in our *Tmu*^+^ mice following DXR treatment. We generated organoids from wild type mice and, after allowing them to establish for 48 h, added 100 ng/mL of either IL-5 or IL-13 to the organoid media. After 4 days of cytokine pre-treatment, the organoids were then treated with DXR for 4 hours and whole mount stained for CC3 to assess apoptosis (Fig 6A). All DXR-treated organoids exhibited an increase in apoptosis, but pre-treatment with IL-5 or IL-13 did not lead to higher rates of DXR-induced apoptosis (Fig. 6B). Interestingly, IL-13 pre-treatment resulted in lower rates of DXR-induced apoptosis. This result suggests that the elevated type 2 cytokines are not driving the increased sensitivity to DXR displayed *in vivo* when *Tmu* is present, and that IL-13 may even confer protection against DXR-induced apoptosis.

**Figure 6:**
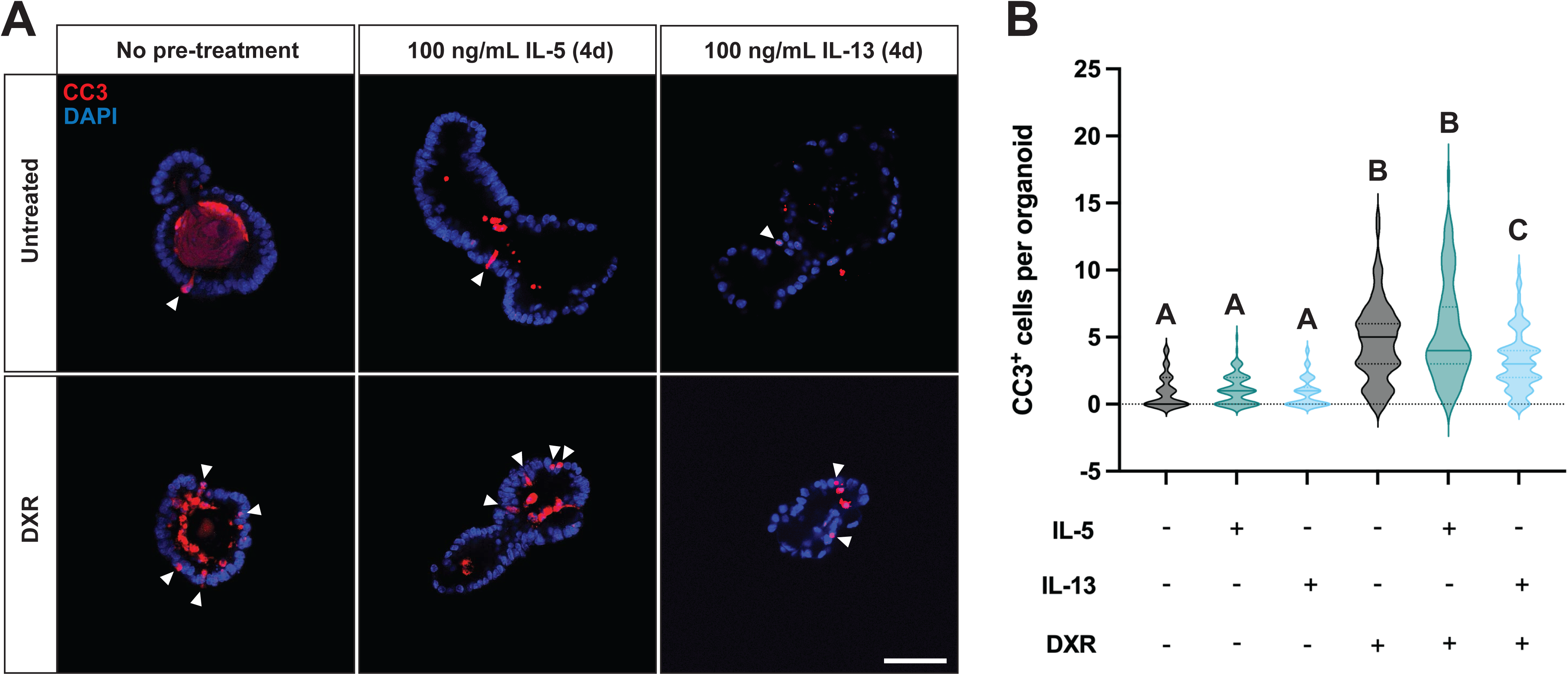
Type 2 cytokines are insufficient to drive increase in DXR-induced apoptosis *in vitro*. (A) Representative images of organoids whole-mount stained for cleaved caspase 3 (CC3) demonstrate the increase in CC3^+^ cells per organoid following DXR treatment. (B) Quantification of CC3^+^ cells per organoid reveals that pre-treating organoids with type 2 cytokines IL-5 or IL-13 does increase the frequency of CC3^+^ cells following DXR, and that organoids pre-treated with IL-13 display fewer CC3^+^ cells following DXR compared to organoids that received no pre-treatment (n = 90 organoids per condition, different letters indicate significant differences, p < 0.0001)

## DISCUSSION

While it is well known that intestinal microbiota influence mucosal biology, the potential contribution of eukaryotes to this interaction has gone largely underappreciated. Recent reports suggest that *Tmu* and related parabasalids are commonly present in the GI tracts of both wild and laboratory mice (8–10, 12, 33, 34). Evidence supports that parabasalids species belonging to the genus Tritrichomonas can be found in the GI tracts of ∼45% of laboratory mice and up to ∼25% of non-industrialized human populations (14, 33). Our findings corroborate previous reports that *Tmu* drives a type 2 innate immune response accompanied by tuft cell hyperplasia, and we contribute a new understanding of how the presence of this often overlooked commensal can alter the intestinal epithelial response to DXR.

Eukaryotic organisms are natural members of the human microbiome, as well, ranging from true parasites to beneficial symbionts. Eukaryome diversity tends to be higher in non-industrialized populations, among which *Tmu* and other *Tritrichomonas* species have been reported (14, 35, 36). Close trichomonad relatives of *Tmu* are also known to colonize the human GI tract, including *Pentatrichomonas hominis* and *Dientamoeba fragilis* (37). *D. fragilis* is widely reported among humans (with a frequency of ∼10%) and, like *Tmu*, largely colonizes healthy individuals asymptomatically (34, 35). Humans also share the same tuft-ILC2 pathway that has been described in mice (38). For these reasons, it is important to understand how these eukaryotic protists influence intestinal biology and drug response. It is possible that the severity of GI side effects from DXR administration is influenced by eukaryotic organisms that colonize the GI tract, and may be of greater concern in non-industrialized populations.

Our findings conflict with the existing model for how the intestinal epithelium responds to DXR. We chose to evaluate tissue at 120 h post-DXR because it had been documented that villus height, crypt depth and density, proliferation, *Lgr5* expression, and OLFM4 expression are all significantly impacted at this timepoint (4, 6, 7, 25). However, we did not see any change in these metrics in our control mice in response to DXR, and only saw reductions in proliferation and *Lgr5* and OLFM4 expression in our *Tmu*^+^ mice. It is worth noting that several of these previous studies found that some of these responses to DXR are dependent upon the presence of intestinal bacteria, reporting findings more closely resembling ours in germ-free mice or mice treated pre-treated with antibiotics (6, 7). This might suggest that the conventional mice used in studies demonstrating severe post-DXR phenotypes were host to an undetected microbe that elicits secondary inflammatory response. Notably, our results are more consistent with those from a recent study performed in rats, which also reports no significant weight loss, no change in crypt depth, and minimal loss of villus height in response to DXR (5). This highlights the important of understanding the influence of distinct microbes on the intestinal response to DXR, which evidently significantly impact the extent of damage caused by the drug.

We do confirm a spike of apoptosis at 6 h post-DXR, which was previously shown to be independent of the microbiome (7). Further, we find that a higher rate of DXR-induced apoptosis occurs in mice colonized with *Tmu*. We also confirm a loss of DCLK1 in response to DXR in our control mice, which was previously reported (4). It was surprising, however, that DCLK1 was upregulated in response to DXR in our mice colonized with *Tmu*. While we demonstrated that this response was not being driven by upregulation of canonical tuft cell-promoting transcription factor expression, it is possible that *Dclk1* transcript and protein expression is being upregulated following DXR due to increased activity of existing transcription factors. Additionally, it was surprising that previously reported fISC transcripts *Ly6a* and *Clu* were not significantly upregulated in response to DXR injury in our mice. It could be the case that expression of these transcripts are not significantly changed at the specific timepoints we evaluated.

We also corroborate that *Tmu* is associated with a type 2 innate immune response in the small intestine, though the cytokines produced by this immune response are not sufficient to drive the increase in DXR-induced apoptosis. It is possible that this increased apoptotic response to DXR is being driven by some other secondary effect of *Tmu* on the intestinal microenvironment or by the inflammatory and immune response it elicits. Ultimately, these findings emphasize the potential impact of intestinal microbes, including those not routinely screened for, on intestinal biology and drug response.

## SUPPLEMENTAL MATERIAL

Supplemental Table S1

## ACKNOWLEDGEMENTS

We thank Drs. Roger Deal, Ken Moberg, Chris Scharer, Bing Yao, and members of the Gracz Lab for constructive discussions and critical reading of the manuscript. We also thank Dr. Peijan He for use of his tissue homogenizer, Dr. Brian Robinson for use of his color camera microscope to obtain images of H&E stained tissue sections, and Dr. Scott Magness for providing additional wild-type mice for experiments.

## AUTHOR CONTRIBUTIONS

NVJ, ARG, and ADG conceived and designed the experiments. NVJ, ARG, SS, and GA performed experiments and analyzed data. NVJ interpreted results of experiments, prepared figures, and drafted manuscript. NVJ and ADG edited and revised manuscript. ADG approved final version of manuscript.

